# Imaging neural activity in the ventral nerve cord of behaving adult *Drosophila*

**DOI:** 10.1101/250118

**Authors:** Chin-Lin Chen, Laura Hermans, Meera C. Viswanathan, Denis Fortun, Michael Unser, Anthony Cammarato, Michael H. Dickinson, Pavan Ramdya

**Affiliations:** Brain Mind Institute, École Polytechnique Fédérale de Lausanne, Lausanne, CH-1015, Switzerland; Interfaculty Institute of Bioengineering, École Polytechnique Fédérale de Lausanne, Lausanne, CH-1015, Switzerland; Department of Medicine, Johns Hopkins University, Baltimore, MD 21205, USA; Department of Physiology, Johns Hopkins University, Baltimore, MD 21205, USA; Biomedical Imaging Group, École Polytechnique Fédérale de Lausanne, Lausanne, CH-1015, Switzerland; Signal Processing core of the Center for Biomedical Imaging (CIBM-SP), École Polytechnique Fédérale de Lausanne, Lausanne, CH-1015, Switzerland; Biology and Biological Engineering, California Institute of Technology, Pasadena, CA 91125, USA

## Abstract

To understand neural circuits that control limbs, one must measure their activity during behavior. Until now this goal has been challenging, because the portion of the nervous system that contains limb premotor and motor circuits is largely inaccessible to large-scale recording techniques in intact, moving animals – a constraint that is true for both vertebrate and invertebrate models. Here, we introduce a method for 2-photon functional imaging from the ventral nerve cord of behaving adult *Drosophila melanogaster*. We use this method to reveal patterns of activity across nerve cord populations during grooming and walking and to uncover the functional encoding of moonwalker ascending neurons (MANs), moonwalker descending neurons (MDNs), and a novel class of locomotion-associated descending neurons. This new approach enables the direct investigation of circuits associated with complex limb movements.

## Introduction

Limbs allow animals to rapidly navigate complex terrain, groom, manipulate objects, and communicate. In vertebrates, neural circuits in the spinal cord coordinate the actions of each arm or leg. Circuits within the thorax perform comparable tasks in insects. The thoracic segments of the fruit fly, *Drosophila melanogaster,* house the ventral nerve cord (VNC) which is a fusion of three thoracic and eight abdominal ganglia. The VNC contains six spherical neuromeres, each controlling one leg, a flat dorsal neuropil associated with the neck, wing, and halteres, and a set of intermediate neuropils including the tectulum that may coordinate the action of the legs and wings. Also within the thorax are descending and ascending axons that connect the VNC to the brain run within a pair of neck or cervical connectives, which – like the VNC – are inaccessible in most preparations.

In recent years, the VNC of adult *Drosophila* has gained attention as the site where some higher-order decisions are transformed into actions. Adult flies engage in complex limbed behaviors including walking^1,2^, reaching^3^, escape jumping^4^, courtship tapping^5^, aggressive boxing^6^, and grooming^7^. Our current understanding of how the VNC coordinates these actions is entirely based on behavioral genetics or recordings from a few neurons in tissue explants^8^, immobilized animals^9-11^, or sharp electrode studies in larger insects^12,13^.

To fully understand how VNC circuits orchestrate limb movements, it is necessary to record the activity of individual cells and populations of neurons during behavior. To date, these experiments have not been performed in *Drosophila* due to the difficulty of accessing the VNC in intact, behaving animals. Here we describe a preparation that overcomes this obstacle and makes it possible to record VNC population dynamics in adult animals during walking, grooming, and other actions involving limb movement.

## Results

The VNC lies on the thoracic sternum – a cuticular structure that anchors the leg muscles and the proximal leg segments to the thorax (**Fig. 1a**). Consequently, it is difficult to access the VNC by removing ventral thoracic cuticle without destroying musculoskeletal elements required for limb movement. We chose instead to access the VNC dorsally at the expense of flight-related behaviors^14^. This approach requires removing the prescutum and scutum of the dorsal thoracic cuticle, the indirect flight muscles (IFMs), and transecting the proventriculus, crop, and salivary glands of the gut (**Fig. 1a**, see Methods).

**Figure 1.**
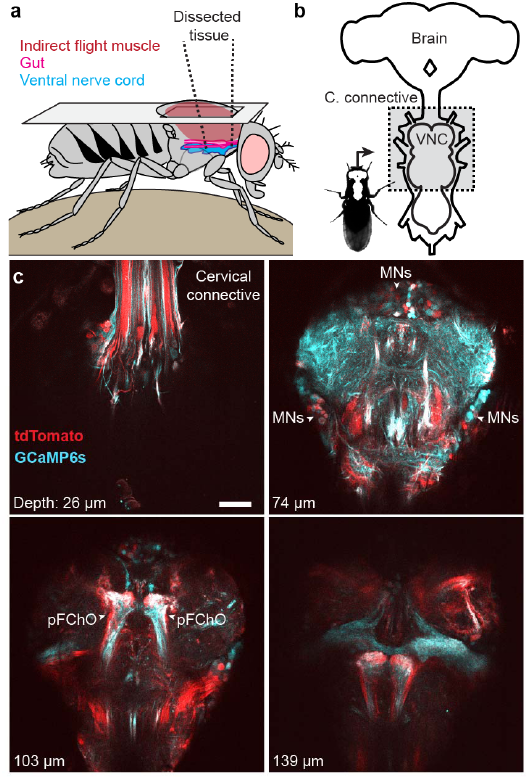
Dissection for imaging the adult *Drosophila* ventral nerve cord (VNC). **(a)** Schematic of the dorsal thoracic dissection. **(b)** Overview of newly accessible nervous tissue following the thoracic dissection. **(c)** Horizontal sections of the VNC imaged at different depths in an animal expressing GCaMP6s (cyan) and tdTomato (red) throughout the nervous system (*GMR57C10>GCaMP6s; tdTomato*). Motor neurons (MNs) and prothoracic (pFChO) femoral chordotonal organs are indicated by white arrowheads. Scale bar is 30 μm.

Using this technique, it is possible to uncover the VNC for functional imaging in flies that are still capable of exhibiting robust behavior, such as walking and grooming, for at least 2-4 hours. To illustrate the extent of VNC access, we drove expression of the genetically encoded calcium indicator, GCaMP6s^15^, together with a fiduciary fluorophore, tdTomato^16^, throughout the entire nervous system (*GMR57C10>GCaMP6s; tdTomato*)^17^, (**Fig. 1b-c** and **Supplementary Video 1**). To perform 2-photon microscopy in semi-intact, behaving animals, we constructed a customized fly holder and spherical treadmill (**Supplementary Fig. 1a**) that, in contrast to previous methods used to record neural activity in the brain^14,18,19^, permits unobscured optical access to the VNC and videography of leg movements (**Supplementary Fig. 1b**).

By focusing on dorsoventral horizontal image planes in animals expressing GCaMP6s and tdTomato pan-neuronally (*GMR57C10>GCaMP6s; tdTomato*), we could record the detailed time course of neural activity in the right and left prothoracic leg neuromeres during walking and grooming (**Fig. 2a-b,e** and **Supplementary Video 2**). We identified two regions-of-interest (ROIs) in the right prothoracic neuromere that correlated with spontaneous prothoracic leg grooming and walking, respectively (**Fig. 2e**). Alternatively, we could use a piezo-driven microscope objective to acquire coronal x-z image planes. These coronal sections allowed us to simultaneously record activity at different depths of the VNC corresponding to layers housing sensory neuron axons ^20^, interneurons^11^, and motor neuron dendrites^21^ (Fig. 2a,c; Supplementary Video 3), or monitor activity patterns across populations of descending^22,23^ and ascending fibers^8,20,23^ within the cervical connective (**Fig. 2a,d** and **Supplementary Video 4**). Thus, we confirmed that our new preparation provides optical access to previously inaccessible thoracic neural populations in behaving animals.

**Figure 2.**
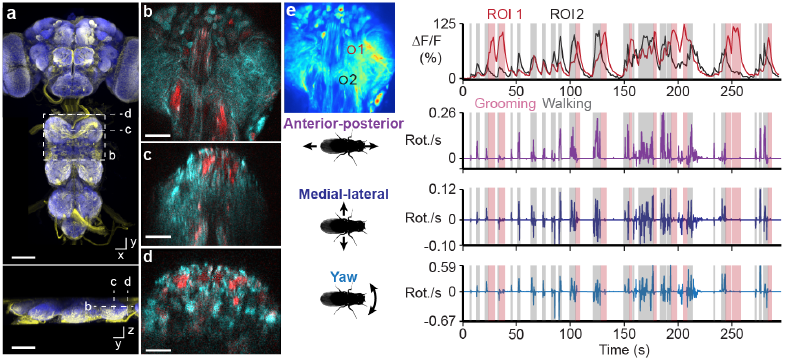
Recording populations of neurons in the VNC during behavior. **(a)** Confocal image of pan-neuronal driver line expression in the brain and VNC. Scale bars are 90 μm. Neuronal GFP (yellow) and neuropil (nc82, blue) are labelled. Dashed lines highlight the imaging modalities made possible by thoracic dissection: **(b)** horizontal section imaging of the VNC (scale bar is 35 μm), **(c)** coronal section imaging of the VNC (scale bar is 50 μm), and **(d)** coronal section imaging of the cervical connective (scale bar is 35μm). All three modalities are illustrated by imaging flies expressing GCaMP6s and tdTomato throughout the nervous system (*GMR57C10>GCaMP6s; tdTomato*). **(e)** %ΔF/F image of a VNC horizontal section of the same animal in **(b)**. ROI-associated fluorescence signals during walking (gray) and grooming (pink) are shown on the top-right. Corresponding rotations of the spherical treadmill are shown on the bottom-right.

Using *Drosophila,* it is possible to repeatedly and systematically investigate the functional properties of sparse sets of genetically-identifiable neurons. In one recent study, a thermogenetic activation screen was used to identify a pair of descending neurons – Moonwalker Descending Neurons (MDNs) – which cause flies to walk backwards^23^. Additionally, concurrent thermogenetic activation of a set of ascending neurons that project from the VNC to the brain – Moonwalker Ascending Neurons (MANs) – resulted in even more sustained backwards walking, perhaps by arresting forward walking^23^. While these activation experiments show that these neurons play an important role in the control of backwards walking, their native activity patterns and the means by which they regulate and report limb movements remain unknown.

Because MAN and MDN axons terminate in the gnathal ganglia (GNG) and the VNC, which are both relatively inaccessible regions of the nervous system, it is difficult to record the activity of these cells during any behavior. We used our functional imaging approach to overcome this challenge and recorded the activity of this set of ascending and descending interneurons within the VNC. Because of the vertical movement artifacts associated with walking, we imaged the activity of MAN axons through coronal sections within the cervical connective (**Fig. 3a**). With this approach, axons are visible as small ellipses (**Fig. 3b**). The *MAN* split*-*GAL4 line we used drives expression in a pair of dorsal and a pair of ventral neurons. We focused our analysis on the dorsal pair of neurons – hereafter referred to as dMANs – because they showed conspicuous changes in activity (**Fig. 3c**). The activity of left and right dMANs were strongly correlated (**Supplementary Fig. 2a**; Pearson’s r = 0.96 ± 0.01, n = 5 flies), allowing us to study their collective response properties. Specifically, we automatically identified the occurrence of transient increases in dMAN fluorescence – referred to as ‘events’ – and examined the corresponding behavioral changes reflected in the spherical treadmill data (see Methods). Our analysis revealed that dMAN events were associated with rapid bimodal anterior-posterior rotations of the spherical treadmill (**Fig. 3d**, n = 746 left and 748 right dMAN events from 9773 s of data from 5 flies). By close inspection of the video data, we observed that these rotations occur when flies extend all six legs to push down on the ball (**Supplementary Videos 5** and **6**).

**Figure 3.**
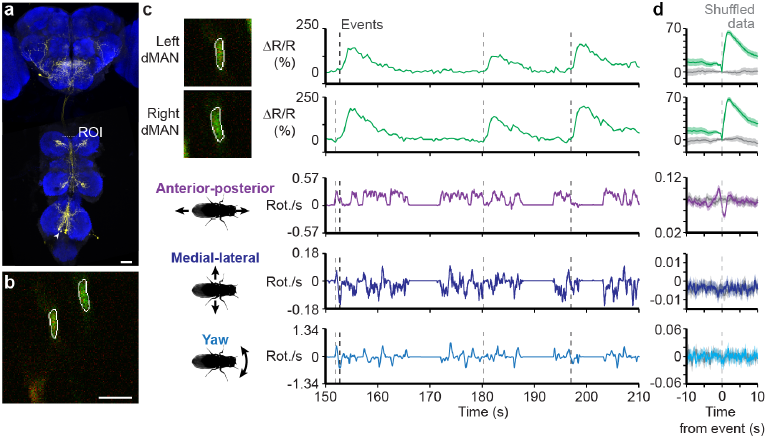
Recording the activity of dorsal Moonwalker Ascending Neurons (dMANs) during behavior. **(a)** Confocal image of *MAN-Gal4* driver line expression in the brain and VNC (scale bar is 40 μm). Neuronal GFP (yellow) and neuropil (nc82, blue) are labelled. A dashed white line highlights the x-z plane imaged. **(b)** Coronal section of the thoracic cervical connective in an animal expressing GCaMP6s and tdTomato in MANs (MAN*>GCaMP6s; tdTomato*). Scale bar is 3.5 μm. **(c)** Separated ROIs **(top-left)** and associated fluorescence signals from left and right dMANs (**top-right**). Corresponding rotations of the spherical treadmill are shown on the bottom right. Events are indicated as dashed gray lines. **(d)** Summary of dMAN activity and spherical treadmill rotations with respect to fluorescence events aligned to 0 s (dashed gray line). Control data in which events are time-shuffled are overlaid in grey. Shown are the means (solid line) and bootstrapped 95% confidence intervals (transparencies).

Next, we asked to what extent MDNs are active during periods of backwards walking, a possibility suggested by behavioral responses to thermogenetic^23^ or optogenetic^24^ MDN stimulation. To address this question, we applied our approach of imaging coronal sections of the thoracic cervical connective using MDN driver line flies expressing GCaMP6s and tdTomato (*MDN-1>GCaMP6s; tdTomato*)(**Fig. 4a-b**). As for dMANs, the activity of pairs of MDNs were strongly correlated (**Fig 4c**), allowing us to focus on their collective response properties (**Supplementary Fig. 2b**; Pearson’s r = 0.93 ± 0.001, n = 3 flies). As predicted, MDNs were active prior to anterior rotations of the spherical treadmill, corresponding to brief episodes of backward walking (**Fig. 4c-d**, n = 900 left and 900 right MDN events from 3 flies and 7790 s of data; **Supplementary Videos 7** and **8**).

**Figure 4.**
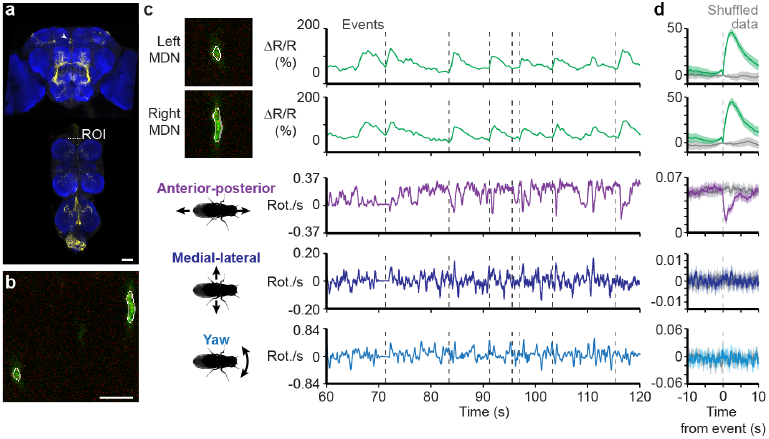
Recording the activity of Moonwalker Descending Neurons (MDNs) during behavior. **(a)** Confocal image of *MDN-1-Gal4* driver line expression in the brain and VNC (scale bar is 40 μm). Neuronal GFP (yellow) and neuropil (nc82, blue) are labelled. A dashed white line highlights the x-z plane imaged. **(b)** Coronal section of the thoracic cervical connective in an animal expressing GCaMP6s and tdTomato in Moonwalker Descending Neurons (MDN-1*>GCaMP6s; tdTomato*). Scale bar is 6 μm. **(c)** Separated ROIs **(top-left)** and associated fluorescence signals from left and right MDNs (**top-right**). Corresponding rotations of the spherical treadmill are shown on the bottom right. Events are indicated as dashed gray lines. **(d)** Summary of MDN activity and spherical treadmill rotations with respect to fluorescence events aligned to 0 s (dashed gray line). Control data in which events are time-shuffled are overlaid in grey. Shown are the means (solid line) and bootstrapped 95% confidence intervals (transparencies).

In addition to resolving the functional properties of previously identified neurons, our method facilitates the discovery of novel cell classes that are active during walking, grooming, and other behaviors involving the limbs or abdomen. As a proof-of-concept, we selected four split-GAL4 lines^25^ that drive sparse expression in pairs of descending neurons^22^ whose axons project to leg neuromeres in the VNC (classes DNa01, DNb06, DNg10, and DNg13). Among these, we found that DNa01 neurons – hereon referred to as A1 cells – were active in a manner that was clearly linked to locomotor state (*A1>GCaMP6s; tdTomato*)(**Fig. 5a-b** and **Supplementary Video 9**). The activity of left and right A1 neurons were not highly correlated (**Fig. 5c** and **Supplementary Fig. 2c**; Pearson’s r = 0.53 ± 0.17, n = 4 flies). Therefore, we investigated the response properties of the left and right cells separately. We found that although the activities of both cells are linked to forward walking, events associated only with left A1 activity were correlated with positive medial-lateral and yaw rotations, or rightward turning by the fly (**Fig. 5d** and **Supplementary Video 10**; n = 1644 events from 4 flies and 8784 s of data). As expected from bilateral symmetry, activity in the right A1 neuron coincided with negative medial-lateral and yaw rotations, or leftward turning (**Fig. 5e** and **Supplementary Video 11**; n = 1651 events from 4 flies and 8784 s of data).

**Figure 5.**
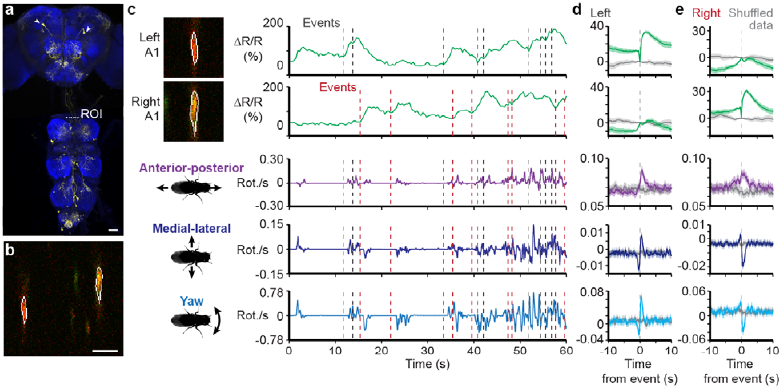
Recording the activity of A1 neurons during behavior. **(a)** Confocal image of *DN0a1-Gal4* driver line expression in the brain and VNC (scale bar is 40 μm). Neuronal GFP (yellow) and neuropil (nc82, blue) are labelled. A dashed white line highlights the x-z plane imaged. **(b)** Coronal section of the thoracic cervical connective in an animal expressing GCaMP6s and tdTomato in A1 neurons (A1*>GCaMP6s; tdTomato*). Scale bar is 5 μm. **(c)** Separated ROIs **(top-left)** and associated fluorescence signals from left and right A1 neurons (**top-right**). Corresponding rotations of the spherical treadmill are shown on the bottom-right. Events are indicated as dashed gray and red lines for left and right A1 neuron events, respectively. **(d)** Summary of A1 neural activity and spherical treadmill rotations with respect to left A1 neuron fluorescence events aligned to 0 s (dashed gray line). **(e)** Summary of A1 neural activity and spherical treadmill rotations with respect to right A1 neuron fluorescence events aligned to 0 s (dashed gray line). Control data in which events are time-shuffled are overlaid in grey. Shown are the means (solid line) and bootstrapped 95% confidence intervals (transparencies).

This approach for recording neural activity in the VNC of behaving *Drosophila* opens up many new avenues for studying premotor and motor circuits. Nevertheless, we can imagine further improvements that will accelerate the study of the thoracic nervous system. For example, in our preparation we found it challenging and time-consuming to remove the indirect flight muscles (IFMs) which fill most of the thorax. Although large, these muscles are quite fragile and tend to disintegrate over the course of an hour after the cuticle of the notum is removed. To increase the efficiency of our dissection, we devised a transgenic strategy to selectively ablate IFMs. We drove the expression of Reaper – a protein promoting apoptosis^26^ – in the IFMs by using a 5’ *Act88F* promotor sequence^27^. *Act88F:Rpr* animals show a nearly complete loss of the IFMs after 7 days post-eclosion (dpe) when raised at 25°C (**Fig. 6a-b**). This loss results in highly elevated or slightly depressed wings, phenotypes identical to those seen in IFM developmental mutants^28^. The heterozygous *Act88F:Rpr* transgenic background greatly accelerated the dorsal thoracic dissection. Although the imaging data in this manuscript were performed without the *Act88F:Rpr* transgene, we envisage that this genetic reagent will greatly simplify and accelerate use of this method in the neuroscience community.

**Figure 6.**
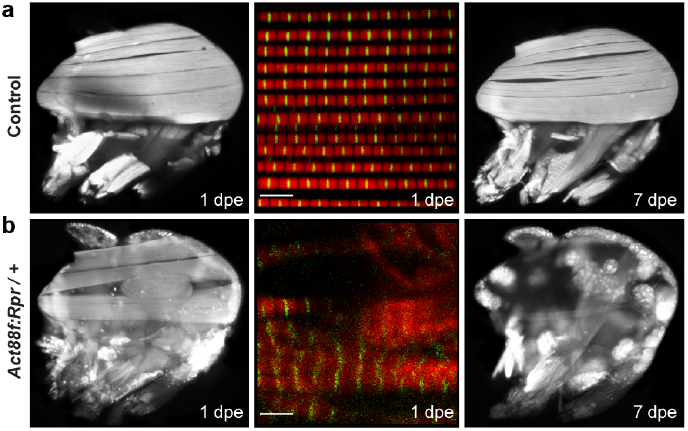
Indirect flight muscles in control and *Act88F:Rpr* animals. Confocal images of dorsal longitudinal IFMs (DLMs) stained with TRITC-phalloidin at 1 dpe **(left)**, 7 dpe **(right)**, or wholemount confocal micrographs of myofibrillar structure **(middle)** for **(a)** wild-type, or **(b)** *Act88F:Rpr* heterozygous flies. Scale bars are 5 μm.

Here we have described a new preparation that enables the visualization of genetically identified neurons in the VNC and cervical connective of *Drosophila* during behavior. We can record the activity of entire neural populations (**Fig. 2**) or measure the encoding of known (**Figs. 3** and **4**) and novel sparse cell classes (**Fig. 5**). This method fills a critical gap in the study of sensory-motor pathways and serves as a complement to ongoing genetic behavioral screens^23,29,30^ aimed at elucidating how populations of neurons coordinate limb movements and orchestrate a variety of legged behaviors.

## Methods

### *Drosophila* lines

Several lines (*GMR57C10-Gal4*, *UAS-GCaMP6s*, *UAS-GCaMP6f*, *UAS-CD4:tdGFP*, and *UAS-tdTomato)* were obtained from the Bloomington Stock Center. *MAN-Gal4* (*VT50660-AD; VT14014-DBD*) and *MDN-1-Gal4* (*VT44845-DBD; VT50660-AD*) were provided by B. Dickson (Janelia Research Campus). *DNa01-Gal4* (SS00731: *GMR22C05-AD; GMR56G08-DBD*), *DNb06-Gal4* (SS02631: *GMR20C04-AD; BJD113E03-DBD*), *DNg13-Gal4* (SS02538: *BJD118A10-AD; BJD123E03-DBD*), and *DNg16-Gal4* (SS01543: *BJD104A07-AD; BJD107F12-DBD*) were provided by G. Rubin (Janelia Research Campus). Transgenic *Actin88F:Rpr* strains (*Act88F:Rpr* flies) were generated using an *Actin88F:eGFP* construct described previously^27^ and injected (BestGene, Chino Hills, CA, USA) with the phiC31-based integration system using the attP18 (X chromosome) landing site^31^.

### Fluorescence imaging of indirect flight muscles

Fluorescent microscopy of hemi-thoraces was performed as described previously^32,33^. Briefly, flies were anesthetized and their heads and abdomens were then removed. Thoraces were fixed overnight in 4% paraformaldehyde at 4°C and rinsed in 1x phosphate buffered saline (PBS) the following day. The specimens were arranged on a glass slide, snap frozen in liquid nitrogen and bisected down the midsagittal plane using a razor blade. IFMs were stained with Alexa-Fluor 568 Phalloidin (1:100 in PBS with 0.1% Triton-X (PBST)) overnight at 4°C, rinsed with PBS and visualized using EVOS^®^ FL Cell Imaging System (Life Technologies) at 4X magnification. For whole-mount imaging of IFM myofibrils, flies were prepared and thoraces bisected as described above. Hemi-thoraces were stained with Alexa-Fluor 568 phalloidin (1:100 in PBST) overnight at 4°C. Samples were rinsed in PBS, mounted with Vectashield (Vector Laboratories) and visualized using a Leica TCS SPE RGBV confocal microscope (Leica Microsystems) at 100X magnification.

### Immunofluorescence imaging of whole-mount brains and ventral nerve cords

Brains and ventral nerve cords (VNCs) were dissected out of 2-3 dpe female flies in PBS. Tissues were then fixed for 20 minutes in 4% paraformaldehyde in PBS at room temperature. After fixation, brains and VNCs were washed 2-3 times in PBS with 1% Triton-X-100 (PBST) and then incubated at 4°C overnight in PBST. Samples were then placed in PBST with 5% normal goat serum (PBSTS) for 20 min at room temperature. They were then incubated with primary antibodies (rabbit anti-GFP at 1:500, Thermofisher; mouse anti-Bruchpilot/nc82 at 1:20, Developmental Studies Hybridoma Bank) diluted in PBSTS for 48 h at 4°C. Brains and VNCs were rinsed 2-3 times in PBST for 10 min each before incubation with secondary antibodies (goat anti-rabbit secondary antibody conjugated with Alexa 488 at 1:500; Thermofisher; goat anti-mouse secondary antibody conjugated with Alexa 633 at 1:500; Thermofisher) diluted in PBSTS for 48 h at 4 °C. Finally, brains and VNCs were rinsed 2-3 times for 10 min each in PBST and mounted onto slides with bridge coverslips in Slowfade mounting-media (Thermofisher).

Samples were imaged using a Carl Zeiss LSM 700 Laser Scanning Confocal Microscope with the following settings: 20X magnification, 8-bit dynamic range, 2x image averaging, 0.52 × 0.52 μm pixel size, 0.57 μm z-axis interval. Standard deviation z-projections of imaging volumes were made using Fiji^34^.

### Thoracic dissection for ventral nerve cord imaging

Custom holders used to mount flies during imaging were fabricated as described previously^35^. For VNC imaging, these stages were modified to have (i) flat rather than folded frames, and (ii) chamfered vertices to make the spherical treadmill visible to optic flow sensors (Shapeways, https://www.shapeways.com/model/upload-and-buy/5963553).

All experiments were performed on 1-3 dpe female flies raised at 25°C on standard cornmeal food on a 12 h light:12 h dark cycle. Flies were anaesthetized at 4°C. A female fly was selected and, in some cases, its wings were clipped to simplify the mounting process. The fly’s dorsal thorax was then pushed through a hole in the steel shim of a custom imaging stage. The stage was then flipped over, UV-curing glue (Bondic, Aurora, ON Canada) was carefully applied around the perimeter of the thorax and hardened through UV illumination (LED-200, Electro-Lite Co. Bethel, CT USA). UV glue was then similarly applied to fix the head and abdomen to the underside of the stage. The stage was then filled with extracellular saline as described previously^14^. Under a high-magnification dissection microscope (Leica M165C), a hypodermic needle (30G, BD PrecisionGlide, Franklin Lakes, NJ USA) was used to slice and lift the cuticle off of the dorsal thorax^36^, being careful not to sever the neck connective. Subsequently, in non-*Act88F:Rpr* animals, a pair of dull forceps was used to remove IFMs, predominantly from the anteriomedial region of the thorax overlying the proventriculus (this step is unnecessary in aged *Act88F:Rpr* animals). This process exposes the dorsal surface of the proventriculus – a large bulbous structure in the gut. With great care, a pair of super-fine forceps was used to grasp and lift the proventriculus to displace much of the gut (including the crop and salivary glands) from the more ventrally located nervous system. With the gut thus elevated, ultra-fine clipper scissors (Fine Science Tools, Foster City, CA USA) were used to transect it at its anterior-most section. The proventriculus was then peeled back and a posterior incision was made to completely remove these portions of the gut, fully revealing the underlying nervous tissue. In some cases, we observed that gut or muscle tissue would begin to obscure the VNC during imaging. Therefore, loose tissue should be removed at this stage while taking great care not to sever the VNC. After each dissection, we examined the extent to which the animal moved its legs in response to a puff of air. This proved to be an accurate predictor of the success of the preparation.

### 2-photon microscopy during behavior

Experiments were performed in the evening zeitgeber time (Z.T.) and animals were typically imaged 30-60 min following dissection. We found that individual animals provided useful data for 2-4 h. Fly holders were secured to a platform raised over the spherical treadmill (**Supplementary Fig. 2a**). The VNC was located using microscope oculars and then aligned to the center of the field-of-view using 2-photon microscopy.

The spherical treadmill was an aluminum rod with a ball-shaped hole milled near its tip^18^. We fabricated 10 mm diameter foam balls (Last-A-Foam FR-7106, General Plastics, Burlington Way, WA USA) and manually spotted them using a Rapidograph pen (Koh-I-Noor, Leeds, MA USA) to provide high-contrast features for optic flow measurements. A 500-600 mL/min stream of filtered and humidified air was passed through the holder using a digital flow controller (Sierra Instruments, Monterey, CA USA). Movements of the ball were measured using optical flow sensors (ADNS3080) outfitted with zoom lenses (Computar MLM3X-MP, Cary, NC USA). The ball and fly were illuminated using a pair of IR LEDs (850-nm peak wavelength) coupled to optic fibers and collimator lenses (ThorLabs, Newton, NJ USA). Optic flow measurements were passed to a microcontroller board (Arduino Mega2560) to be recorded using custom-written Python code. Simultaneously, videography of behaviors on the ball were made using an IR-sensitive firewire camera (Basler, Ahrensburg, Germany) at approximately 30 frames per second.

We performed 2-photon microscopy using a Bergamo II microscope (ThorLabs) outfitted with two GaAsP PMT detectors for GCaMP6 and tdTomato imaging, respectively, and coupled to a Ti:Sapphire laser (MaiTai DeepSee, Newport Spectra-Physics, Santa Clara, CA USA) tuned to 930 nm. We used an Olympus 20X objective water-immersion lens with 1.0 NA (Olympus, Center Valley, PA USA). The microscope was controlled using ThorImage software (ThorLabs). Occasionally, a puff of air was used to elicit walking behaviors. These puffs were digitally encoded (Honeywell AWM 3300V, Morris Plains, NJ USA). Custom ROS software interfaced through an analog output device (Phidgets, Calgary, Canada) to ThorSync software (ThorLabs) was used to synchronize optic flow measurements, behavior videography, air puff measurements, and 2-photon image acquisition. For coronal section imaging, a piezo collar (Physik Instrumente, Karlsruhe, Germany) was used for rapid z-axis movements of the microscope objective lens.

### Data analysis

We analyzed all data using custom scripts written in Python. Because the data acquisition frequency differed for optic flow, behavior videography, and 2-photon imaging we interpolated signals to that of the highest frequency. Subsequently, optic flow data were smoothed using a running average and then translated into rotations s^-1^ for the anterior-posterior, medial-lateral, and yaw axes as described in^18^.

#### Pan-neuronal image registration, ROI identification, and fluorescence processing (related to Fig. 2)

We observed that large tissue deformations could occur during behavior. Therefore, we performed post-hoc registration of pan-neuronal imaging data. To do this, we registered all frames of an imaging experiment with one reference image. Because the complexity of the deformations could not be captured by simple parametric motion models (e.g., affine transformations), we used a non-parametric, variational approach, designed to model arbitrarily complex deformations. We computed the motion field **w** between the reference image, denoted *I*_*r*_, and the image at time *t*, denoted *I*_*t*_, by solving the minimization problem

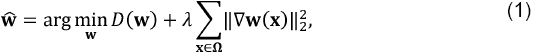

where *D*(**w**) is a data fitting term, the second term is a regularization promoting smoothness of w by penalizing its gradient ∇**w** ^37^, Ω is the discrete image domain, and the parameter *λ* weights the contributions of the two terms.

The sequence tagged with GCaMP6s images is characterized by two main difficulties for motion estimation. First, fast motion of the fly induces very large deformations. Second, the activation of neurons produces large local intensity changes between corresponding pixels in *I*_*r*_ and *I*_*t*_. To address these issues, we defined a data term of the form

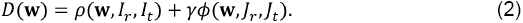

The first term models the standard assumption of conservation of intensity along the trajectory of each pixel. It is defined by

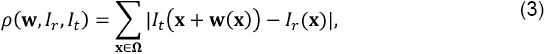

where we use an ℓ_1_ norm to gain partial robustness to intensity changes^38^. The second term in (2) is a feature matching constraint inspired by Revaud and co-workers^39^, written as

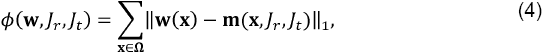

where the images *J*_*r*_ and *J*_*r*_ are the analog of *I*_*r*_ and *I*_*t*_ in the second channel tagged with tdTomato, for which we do not expect activity-dependent intensity changes. Minimizing the function φ favors motion vectors **w**(**x**) to be close to feature correspondences **m**(**x**,*J*_*r*_,*J*_*t*_), computed on a sparse set of relevant keypoints. We obtain **m** with the feature matching algorithm proposed by Revaud and co-workers^39^, which is specifically designed to handle large image deformations. We compute **m** using *J*_*r*_ and *J*_*t*_, such that the correspondences are also insensitive to the intensity changes between *I*_*r*_ and *I*_*t*_. As a result, the estimation is guided by reliable feature matches. We found that it is necessary to keep a standard data term (3) defined on the GCaMP6s imaging channel, because the tdTomato channel may not provide information in some regions of the image. The parameter γ balances the two terms in (2).

We solved the optimization problem (1) with an alternated direction method of multiplier (ADMM) algorithm^40^. We introduced two splitting variables, associated with the regularization and the feature matching terms, respectively. Each sub-problem of the algorithm was solved analytically. We used parts of the inverse problems library described in ^41^. A post processing based on weighted median filtering was applied with the method of ^42^.

From these registered imaging data, regions-of-interest (ROIs) were manually selected. %ΔF/F values were then calculated from fluorescence signals averaged within the ROIs. ΔF = F*_t_* − F, where F*_t_* is the average fluorescence within an ROI at time, *t.* F is a baseline fluorescence signal that was calculated as the average pixel value for the first ten sequential GCaMP6s images for which no cellular activity was observed (i.e., minimal and unchanging GCaMP6s fluorescence).

#### Sparse neuron ROI identification, and fluorescence processing (related to Figs. 3-5)

For single-neuron fluorescence data, ROIs were selected using custom Python scripts relying on OpenCV and Numpy libraries. First, a reference frame was selected for which software identified all potential ROIs. To do this, the GCaMP6s channel image was smoothed to reduce background noise and then an Otsu filter threshold was applied to the image. An erosion factor was then applied on all objects detected within the image.

Contours of all detected objects were then presented to the user for manual selection. Once these reference ROIs were selected for left and right neurons, we used a cross-correlation-based image registration algorithm^43^ to identify the most likely left and right ROIs for each image frame based on those manually selected on the reference frame. A second script was used to manually verify automatically selected ROIs and, if incorrect, to display all potential ROIs within the frame for manual selection. If chosen erosion values yielded malformed ROI shapes, another script was used to manually position elliptical ROIs with arbitrary orientations on a given frame. Finally, binary ROI images were used as an image mask to extract mean fluorescence intensities from the original GCaMP6s or tdTomato images. These signals were reported as %ΔR/R as in ^44^ to reduce the effects of motion artifacts on fluorescence signals. Due to the absence of stimuli for eliciting behaviors, the baseline R was calculated as the minimum ratio of GCaMP6s / tdTomato within a 2.5 s bin.

To detect transient increases in activity above the %ΔR/R baseline, we developed an algorithm based partly on ^45^. We first determined when the first order derivative of the %ΔR/R signal crossed an arbitrary threshold, which was calculated as a percentile determined by examining all derivatives values for a given neuron class (i.e., MDN, MAN, or A1). We reasoned that threshold values should be characteristic and potentially different for each neuron class because fluorescence dynamics are related to intrinsic physiological properties that can differ across neuron classes but not across experiments investigating a single class. We set this threshold for the derivative value as the 97.5^th^ percentile for MDNs and dMANs and 90^th^ percentile for A1 neurons. A lower threshold value was selected for A1 neurons because many more fluorescence transients were observed in A1 %ΔR/R fluorescence traces. These transients would be overlooked using the 97.5^th^ percentile. To identify the onset of fluorescence increases we found the nearest preceding time point where the derivative crossed zero (i.e., typically an inflection between decreases and increases in fluorescence). This zero-crossing is considered the time-point of an ‘event’ associated with the identified fluorescence increase. Events detected close to one another with no intervening derivative zero-crossing were compressed into one event associated with the first time point. There were ∼10 separate experiments per animal. Events in the first and last 10 s of each experiment were not considered since the window for data presentation encompassed 10 s before and 10 s after each event.

Because left and right MDN and dMAN cells strongly covaried (**Supplementary Fig. 2**), an additional step was performed for event detection: if events were detected in both left and right neurons within 2 s of one another, both events were retained; otherwise, an event identified for neuron A (e.g., left MDN) and not neuron B (e.g., right MDN) was also added to neuron B’s event library.

By contrast, left and right A1 neural activities did not strongly covary. Therefore, analyzed events associated uniquely to one and not the other neuron. To accomplish this, if an event was detected in both left and right A1 neurons within a time window of 0.25 s, neither of the events were logged for analysis.

%ΔR/R and optic flow traces linked to each event were aligned by setting the event time points to 0 s. We then computed the mean and bootstrapped 95% confidence intervals for these aligned traces using the Python Seaborn library. Optic flow and %ΔR/R measurements were downsampled to 500 values/s for this analysis. To increase clarity, %ΔR/R traces were baseline-subtracted to make them zero at the time of the event in the summary **Figs. 3d, 4d**, and **5d-e**. Control shuffled data (gray traces) were computed by instead assigning random time-points in place of real, identified events. These random time points were treated as real events and their mean and bootstrapped 95% confidence intervals were computed and plotted for comparison.

#### Covariance analysis (related to Supplementary Figure 2)

Covariance analysis was performed with a custom Python script using Matplotlib and Numpy libraries. Scatter plots were computed comparing left and right neuron %ΔR/R values from all experiments for each fly separately. All data were included for analysis in these scatter plots with the exception of two MDN flies that exhibited unusually low fluorescence values. Pearson’s r values are reported as mean ± standard deviation.

#### Event-related behaviors (related to Supplementary Videos 6, 8, 10, and 11)

Events chosen for behavioral summary videos were chosen from automatically detected events as described above. For dMANs, events were manually selected from those that maximized the difference in anterior-posterior ball rotations between 1 s before the event and 2 s after the event. For MDNs, events were manually selected from among those that minimized anterior-posterior ball rotations up to 2 s after the event. For A1 neurons, events were manually selected from among those that maximized the average yaw ball rotations (positive for left A1 neuron examples and negative for right A1 neuron examples) for up to 2 s after the events.

## Acknowledgments

We thank B.J. Dickson (Janelia Research Campus, VA) for *MDN-1-Gal4* and *MAN-Gal4* fly strains. We thank G. Rubin (Janelia Research Campus, VA) for *DNa01-Gal4*, *DNb06-Gal4*, *DNg13-Gal4*, and *DNg16-Gal4* fly strains. AC acknowledges support from the National Institutes of Health (R01HL124091). MHD acknowledges support from the National Institute of Neurological Disorders and Stroke of the National Institutes of Health (U01NS090514). PR acknowledges support from the Swiss National Science Foundation (31003A_175667).

## Author contributions

C.L.C. generated strains; performed experiments; analyzed data

L.H. analyzed data

M.C.V. performed experiments; analyzed data

D.F. wrote analysis code

M.U. supervised the project

A.C. designed and supervised the project

M.H.D. designed and supervised the project

P.R. conceived of, designed, and supervised the project; performed experiments; analyzed data

All authors contributed to writing the paper

---

**Supplementary Figure 1.**
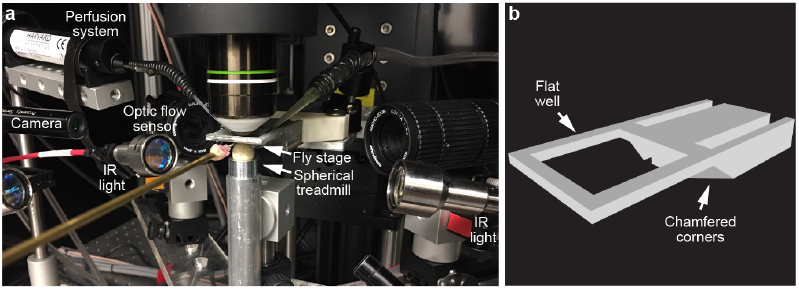
System for VNC imaging. **(a)** Photograph of the spherical treadmill system and **(b)** schematic of the custom fly holder used in this study.

**Supplementary Figure 2.**
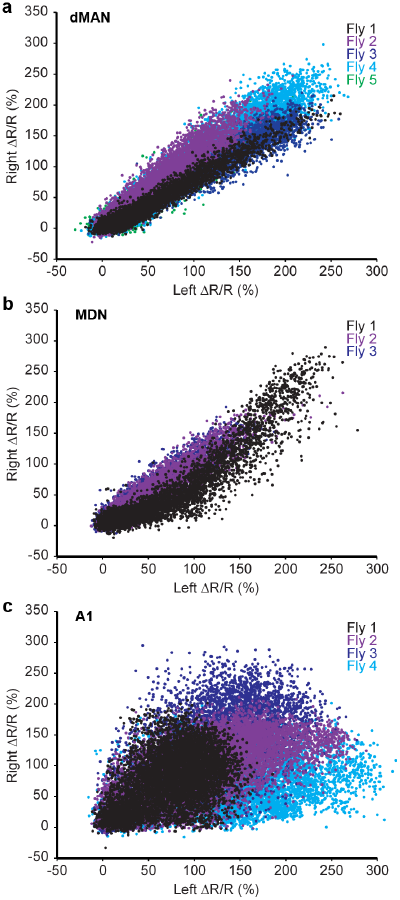
Covariance in fluorescence signals between bilateral pairs of neurons. Scatter plots comparing %ΔR/R signals for left and right **(a)** dMAN, **(b)** MDN, or **(c)** A1 neuron pairs. Data from each animal are color-coded.

## Supplementary Videos

**Supplementary Video 1** | **Extent of VNC imaging volume.** 2-photon imaging of horizontal sections across the dorsal-ventral extent of the VNC and cervical connective. GCaMP6s (cyan) and tdTomato (red) are expressed throughout the nervous system (*GMR57C10>GCaMP6s; tdTomato*). Imaging depth is indicated on the top-left.

**Supplementary Video 2** | **Horizontal VNC imaging.** 2-photon imaging of a single horizontal section of the VNC in a walking and grooming fly. GCaMP6s (cyan) and tdTomato (red) are expressed throughout the nervous system (*GMR57C10>GCaMP6s; tdTomato*). Shown are synchronized raw fluorescence images **(top-left)**, %ΔF/F images **(top-right)**, behavior images **(bottom-left)**, and spherical treadmill rotations along the anterior-posterior (‘AP’), medial-lateral (‘ML’), and yaw axes **(bottom-right)**. Experimenter-administered air puffs are indicated by the appearance of red boxes above behavior video images. Video is 4X faster than real-time.

**Supplementary Video 3** | **Coronal VNC imaging.** 2-photon imaging of a single coronal section of the VNC in a walking fly. GCaMP6s (cyan) and tdTomato (red) are expressed throughout the nervous system (*GMR57C10>GCaMP6s; tdTomato*). Shown are synchronized raw fluorescence images **(top-left)**, %ΔF/F images **(top-right)**, behavior images **(bottom-left)**, and spherical treadmill rotations along the anterior-posterior (‘AP’), medial-lateral (‘ML’), and yaw axes **(bottom-right)**. Video is 4X faster than real-time.

**Supplementary Video 4** | **Coronal cervical connective imaging.** 2-photon imaging of a single coronal section of the cervical connective in a walking fly. GCaMP6s (cyan) and tdTomato (red) are expressed throughout the nervous system (*GMR57C10>GCaMP6s; tdTomato*). Shown are synchronized raw fluorescence images **(top-left)**, %ΔF/F images **(top-right)**, behavior images **(bottom-left)**, and spherical treadmill rotations along the anterior-posterior (‘AP’), medial-lateral (‘ML’), and yaw axes **(bottom-right)**. Video is 4X faster than real-time.

**Supplementary Video 5** | **Coronal cervical connective imaging of dorsal Moonwalker Ascending Neurons.** 2-photon imaging of a single coronal section of the cervical connective in a behaving fly. GCaMP6s (cyan) and tdTomato (red) are expressed in MANs (*MAN>GCaMP6s; tdTomato*). Raw fluorescence images of the left and right dMANs are presented and outlined by ROIs **(top-left)**. These images are used to calculate %ΔR/R traces for each neuron **(top-right)**. Corresponding behavior videography **(bottom-left)** and spherical treadmill rotations along the anterior-posterior (‘AP’), medial-lateral (‘ML’), and yaw axes **(bottom-right)** are shown.

**Supplementary Video 6** | **Behavioral responses associated with dorsal Moonwalker Ascending Neuron activity events.** Three example behaviors (rows) for each of three flies (columns) produced at the onset of dMAN fluorescence events. Red square indicates the time of each fluorescence event (t = 0 s). Video is 3X slower than real-time.

**Supplementary Video 7** | **Coronal cervical connective imaging of Moonwalker Descending Neurons.** 2-photon imaging of a single coronal section of the cervical connective in a behaving fly. GCaMP6s (cyan) and tdTomato (red) are expressed in MDNs (*MDN-1>GCaMP6s; tdTomato*). Raw fluorescence images of the left and right MDNs are presented and outlined by ROIs **(top-left)**. These images are used to calculate %ΔR/R traces for each neuron **(top-right)**. Corresponding behavior videography **(bottom-left)** and spherical treadmill rotations along the anterior-posterior (‘AP’), medial-lateral (‘ML’), and yaw axes **(bottom-right)** are shown.

**Supplementary Video 8** | **Behavioral responses associated with Moonwalker Descending Neuron activity events.** Three example behaviors (rows) for each of three flies (columns) produced at the onset of MDN fluorescence events. Red square indicates the time of each fluorescence event (t = 0 s). Video is 3X slower than real-time.

**Supplementary Video 9** | **Coronal cervical connective imaging of A1 Neurons.** 2-photon imaging of a single coronal section of the cervical connective in a behaving fly. GCaMP6s (cyan) and tdTomato (red) are expressed in A1 neurons (*A1>GCaMP6s; tdTomato*). Raw fluorescence images of the left and right A1 neurons are presented and outlined by ROIs **(top-left)**. These images are used to calculate %ΔR/R traces for each neuron **(top-right)**. Corresponding behavior videography **(bottom-left)** and spherical treadmill rotations along the anterior-posterior (‘AP’), medial-lateral (‘ML’), and yaw axes **(bottom-right)** are shown. Video is 2X faster than real-time.

**Supplementary Video 10** | **Behavioral responses associated with left A1 neuron activity events.** Three example behaviors (rows) for each of three flies (columns) produced at the onset of left A1 neuron fluorescence events. Red square indicates the time of each fluorescence event (t = 0 s). Video is 3X slower than real-time.

**Supplementary Video 11** | **Behavioral responses associated with right A1 neuron activity events.** Three example behaviors (rows) for each of three flies (columns) produced at the onset of right A1 neuron fluorescence events. Red square indicates the time of each fluorescence event (t = 0s). Video is 3X slower than real-time.

